# The *Pseudomonas aeruginosa* phosphodiesterase gene *nbdA* is transcriptionally regulated by RpoS and AmrZ

**DOI:** 10.1101/2021.03.31.437996

**Authors:** Katrin Gerbracht, Susanne Zehner, Nicole Frankenberg-Dinkel

## Abstract

*Pseudomonas aeruginosa* is an opportunistic pathogen causing serious infections in immune compromised persons. These infections are difficult to erase with antibiotics, due to the formation of biofilms. The biofilm lifecycle is regulated by the second messenger molecule c-di-GMP (bis-3,5-cyclic di-guanosine monophosphate). *P. aeruginosa* encodes 40 genes for enzymes presumably involved in the biosynthesis and degradation of c-di-GMP. A tight regulation of expression, subcellular localized function and protein interactions control the activity of these enzymes. In this work we elucidated the transcriptional regulation of the gene encoding the membrane-bound phosphodiesterase NbdA. We previously reported a transcriptional and posttranslational role of nitric oxide (NO) on *nbdA* and its involvement in biofilm dispersal. NO is released from macrophages during infections but can also be produced by *P. aeruginosa* itself during anaerobic denitrification. Recently however, contradictory results about the role of NbdA within NO-induced biofilm dispersal were published. Therefore, the transcriptional regulation of *nbdA* was reevaluated to obtain insights into this discrepancy. Determination of the transcriptional start site of *nbdA* by 5’-RACE and subsequent identification of the promoter region revealed a shortened open reading frame (ORF) in contrast to the annotated one. In addition, putative binding sites for RpoS and AmrZ were discovered in the newly defined promoter region. Employing chromosomally integrated transcriptional *lacZ* reporter gene fusions demonstrated a RpoS-dependent activation and AmrZ repression of *nbdA* transcription. In order to investigate the impact of NO on *nbdA* transcription, conditions mimicking exogenous and endogenous NO were applied. While neither exogenous nor endogenous NO had an influence on *nbdA* promoter activity, deletion of the nitrite reductase gene *nirS* strongly increased *nbdA* transcription independently of its enzymatic activity during denitrification. The latter supports a role of NirS in *P. aeruginosa* apart from its enzymatic function.

**IMPORTANCE:** The opportunistic pathogen *Pseudomonas aeruginosa* possesses a network of genes encoding proteins for the turnover of the second messenger c-di-GMP involved in regulating-among others-the lifestyle switch between planktonic, motile cells and sessile biofilms. Insight into the transcriptional regulation of these genes is important for the understanding of the protein function within the cell. Determination of the transcriptional start site of the phosphodiesterase gene *nbdA* revealed a new promoter region and consequently a shortened open reading frame for the corresponding protein. Binding sites for RpoS and AmrZ were identified *in silico* and confirmed experimentally. Previously reported regulation by nitric oxide was reevaluated and a strong influence of the moonlighting protein NirS identified.

## INTRODUCTION

The opportunistic human pathogen *Pseudomonas aeruginosa* is able to form acute and chronic infections, the latter associated with biofilm formation (1). Within biofilms, bacteria are embedded in a self-produced matrix and are highly protected against the host immune system and antibiotic treatments (2, 3). Therefore, biofilm associated infections are difficult to treat and the *P. aeruginosa* biofilm lifecycle has become a well-studied topic in the last decades. Environmental cues like changes in nutrient availability or the diatomic gas nitric oxide (NO) are able to induce biofilm dispersal by promoting a switch between the sessile and planktonic lifestyle of the bacteria (4–6). In general, the biofilm lifecycle is dependent on the second messenger bis-(3,5)-cyclic diguanosine-monophosphate (c-di-GMP). However, c-di-GMP does not only regulate the biofilm lifecycle, but rather is involved in various bacterial processes e.g., motility, secretion systems, virulence and cell cycle progression (7). The intracellular level of c-di-GMP is dependent on diguanylate cyclases (DGC) that build c-di-GMP from two molecules of GMP and c-di-GMP-specific phosphodiesterases (PDE) that hydrolyze c-di-GMP to either pGpG or GMP (8–12). DGC domains contain a conserved GGDEF motif whereas PDE domains contain either an EAL or HD-GYP motif. Typically, bacteria encode multiple DGC, PDE or tandem enzymes. Adjustment of the intracellular c-di-GMP concentration can be achieved by regulating the production of c-di-GMP modulating proteins on different levels: transcription, post-transcription and post-translation.

On the transcriptional level, control of gene expression by various transcription factors or alternative sigma factors that react to changing environmental or growth conditions allows a temporal separation of redundant PDEs or DGCs within a bacterial cell. For instance, in *E. coli* many DGC or PDE encoding genes are under the control of the alternative sigma factor RpoS (σ^S^) which regulates genes for stationary growth phase or stress responses (13). Post-transcriptionally, RNA-binding proteins like CsrA of *E. coli* or RsmA of *P. aeruginosa* are able to influence the translation of DGCs or PDEs by binding to corresponding mRNAs (14, 15). Functional sequestration of DGCs and PDEs within a cell is achieved on the post-translational level. For example, PDEs with EAL-motif often require dimerization to enable c-di-GMP hydrolysis (16). Binding of GTP to the GGDEF domain of the *P. aeruginosa* RbdA enhances PDE activity of the tandem protein (17). Diguanylate cyclases can be object of product feedback inhibition by binding of c-di-GMP to the I-site of the DGC domain (18). Additionally, PDE and DGC domains are often coupled to sensory domains, which allows stimulation of activity in response to environmental signals. Binding of O_2_ to the heme co-factor of the *E. coli* DosP for example is required for the proteins PDE activity (19). In the case of PA0575 of *P. aeruginosa*, binding of L-arginine to a sensory Venus flytrap (VFT) domain stimulates c-di-GMP degradation (20). Another possibility to avoid functional redundancies of c-di-GMP modulating proteins is introduced by the “fountain model”. It proposes spatial sequestration of particular DCGs and PDEs within a cell and the influence on only a local c-di-GMP pool rather than on the global c-di-GMP concentration (21).

In *P. aeruginosa*, the c-di-GMP modulating network consists of 40 DGC, PDE or tandem proteins which contain both domains (22). One of them is the <u>N</u>O-induced <u>b</u>iofilm <u>d</u>ispersion locus <u>A</u> (NbdA). NbdA is a three domain protein, consisting of the membrane anchored MHYT domain, a diguanylate cyclase domain with a degenerated GGDEF motif and a phosphodiesterase domain with an EAL motif (23). NbdA was shown to be a functional phosphodiesterase, lacking DGC activity (23). The MHYT domain of NbdA is predicted to be a sensory domain for diatomic gases like oxygen, NO or carbon monoxide (CO) (24). In a previous study we observed that a *nbdA* deletion mutant showed glutamate-induced biofilm dispersal, but was unable to disperse in response to nitric oxide. Additionally, we suggested transcriptional regulation of *nbdA* by NO as *nbdA* transcript levels were increased in NO-treated planktonic cells or dispersed cells after NO-induced dispersal when compared to untreated planktonic cells (23). In *P. aeruginosa* NbdA is not the only protein involved in NO-induced biofilm dispersal as deletion mutants of the PDEs *rbdA* and *dipA* both display the same phenotype as Δ*nbdA* (25). However, in a biofilm model on airway epithelial cells it was demonstrated that the deletion of neither *nbdA*, *rbdA* nor *dipA* led to a loss of biofilm dispersal in response to NO (26). These contrary findings underline, that c-di-GMP modulating proteins are tightly regulated in *P. aeruginosa* and changes in environmental conditions might impact expression or activity of those enzymes. Therefore, we decided to reevaluate the transcriptional regulation of *nbdA* to obtain insights into this discrepancy and to gain a better understanding of NbdA’s role within the c-di-GMP network of *P. aeruginosa* PAO1.

## MATERIAL AND METHODS

### Bacterial strains and growth conditions

All strains used in this study are listed in Table 1. If not stated otherwise, bacteria were grown in LB medium at 37 °C. For denitrification conditions, growth medium was supplemented with 50 mM KNO_3_. Pseudomonas isolation agar was used for selection of *P. aeruginosa* after mating procedures. Antibiotics were used in following concentrations: gentamicin, 10 μg ml^-1^ (*E. coli*); 75 μg ml^-1^ (*P. aeruginosa*); tetracycline, 5 μg ml^-1^ (*E. coli*); 100 μg ml^-1^ (*P. aeruginosa*).

**Table 1:**
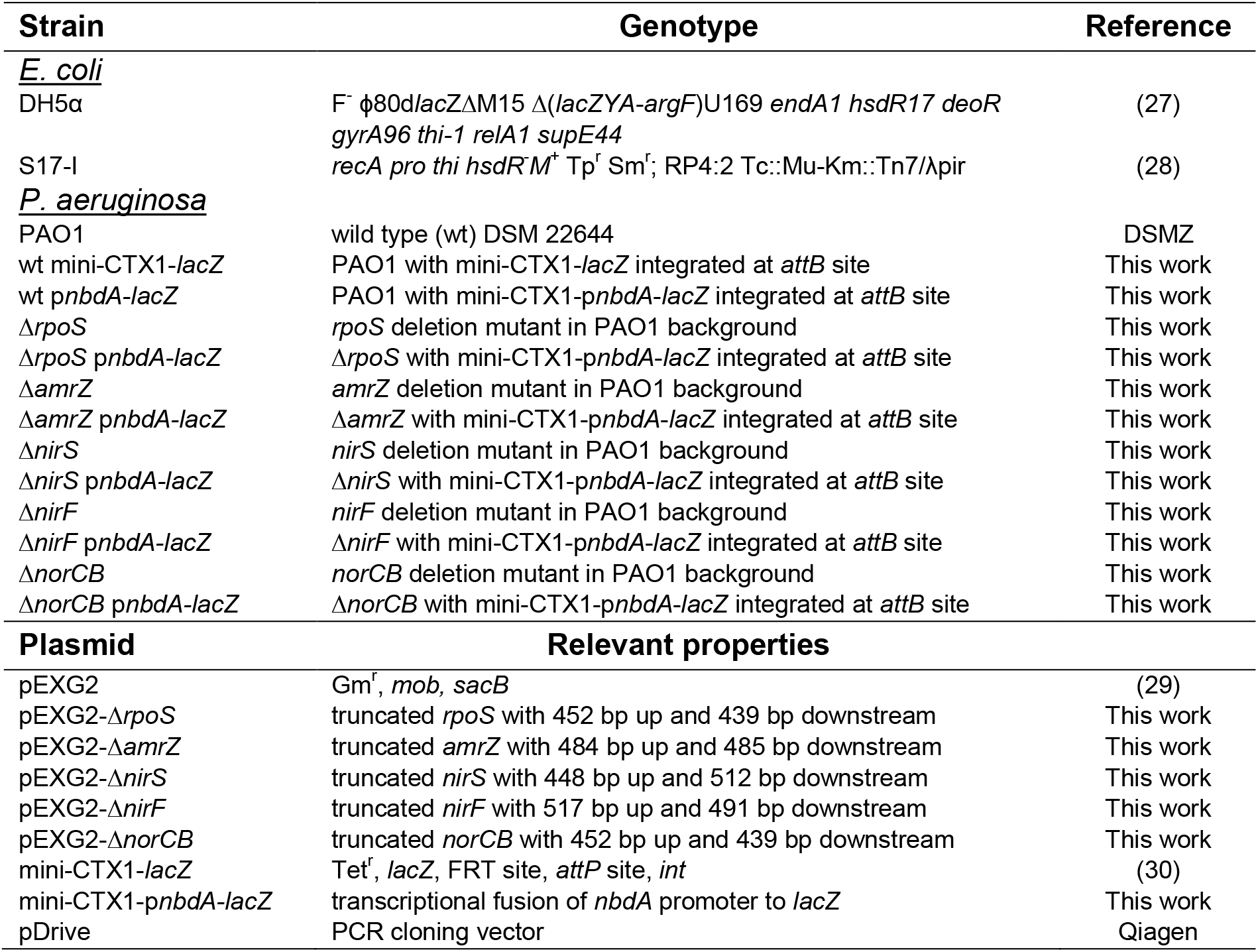
Strains and plasmids used in this study.

### Plasmid and strain construction

Oligonucleotides used for plasmid construction are listed in Table 2. Markerless deletion mutants were produced as described previously with minor modifications (31). DNA fragments were generated for each deletion via splicing-by-overlap extension (SOE) PCR using the corresponding Up and Down primer pairs (Table 2) and integrated into the allelic exchange vector pEXG2 (29). The vector was transferred from *E. coli* S17-I to *P. aeruginosa* PAO1 via biparental mating. Pseudomonas isolation agar supplemented with gentamicin was used to select cells which integrated the allelic exchange vector by homologous recombination. Those cells were streaked twice on LB medium containing sucrose (15 % w/v) to force the second crossover event. Truncation of target genes was verified via colony PCR and sequencing.

**Table 2:**
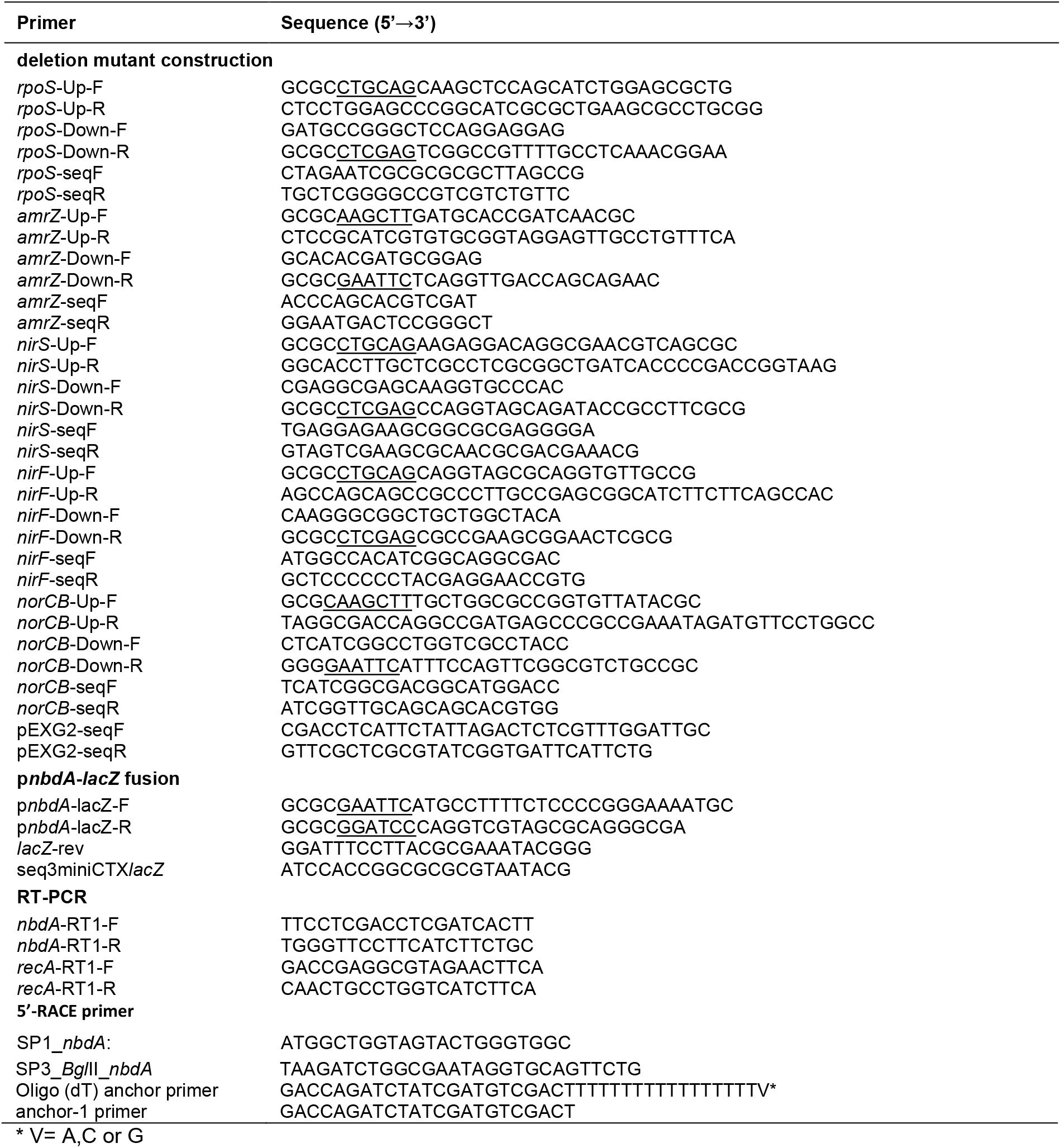
Primers used in this study.

Transcriptional *nbdA-lacZ* fusion was generated in the vector mini-CTX1-*lacZ* (30). A 171 bp fragment of the *nbdA* promoter region and 279 bp of the coding sequence was amplified via PCR and integrated in front of the promoterless *lacZ* gene encoded on the vector. The transcriptional fusion and the empty vector control were transferred to *P. aeruginosa* strains via biparental mating and chromosomally integrated into the *attB* site on the genome via integrase-mediated chromosomal integration.

### β-Galactosidase assay

For measurements of promoter activity the β-galactosidase assay protocol of Miller (32) was modified as follows. Overnight cultures containing the promoter-*lacZ* fusion were diluted to an OD_600_ of 0.01 in 20 ml LB with respective antibiotics. In exponential (4 h) and early stationary growth phase (7 h) OD_600_ was measured and cells of 100 μl culture were harvested by centrifugation. Cells were resuspended in 800 μl Z-buffer (60 mM Na_2_HPO_4_ x 7H_2_O; 40 mM NaH_2_PO_4_ x H_2_O; 10 mM KCl; 1 mM MgSO_4_ x 7H_2_O; 50 mM β-mercaptoethanol). To permeabilize the cells 25 μl of 0.1 % (w/v) sodium-dodecyl sulfate (SDS) and 25 μl chloroform were added. After 5 min incubation 200 μl of 4 mg/ml *ortho-nitrophenyl*-β-D-galactosid (ONPG) were added to the mixture at 37 °C to start the reaction. When β-galactosidase activity was indicated by a color change due to the formation of the yellow colored product *ortho*-nitrophenol, the reaction was stopped by addition of 500 μl 1 M Na_2_CO_3_. Cell debris was precipitated by centrifugation and product formation was measured in the supernatant at OD_420_. The activity of β-galactosidase was calculated as follows: Miller units (MU) = (OD_420_ / (OD_600_ * volume * incubation time)) * 1000.

### RNA extraction and semi quantitative RT-PCR

Bacteria were grown to exponential and early stationary phase as described above. To prevent RNA degradation, cells of an equivalent of OD_600_ = 1 in 1 ml were mixed with 100 μl RNA stop solution (5 % (v/v) phenol in ethanol). Cell pellets were stored at −80 °C or used directly for RNA extraction. RNA was extracted by enzymatic lysis according to Qiagen RNA protect handbook and further purified with the RNeasy Plus Mini kit (Qiagen) following the suppliers instruction. Extracted RNA was transcribed into cDNA using ProtoScript II Reverse Transcriptase (New England Biolabs) according to the manufacturers’ protocol. 2.5 ng of cDNA were used as template for 25 μl semi quantitative RT-PCR reaction. Used primers are listed in Table 2.

### Determination of transcriptional start sites by 5’-RACE

*P. aeruginosa* cells were inoculated 1:100 from an overnight culture in LB medium and incubated 5 h at 37 °C. Total RNA isolation was performed as previously described (33). Primers used for 5’-RACE are listed in Table 2. cDNA was synthesized at 42 °C for 60 min with M-MLV-RT (Promega) using gene-specific primer *SP1_nbdA*. The cDNA was treated with shrimp alkaline phosphatase (New England Biolabs) and purified with MinElute kit (Qiagen). A deoxyadenosine tail was added to the 3’ end of the cDNA using terminal transferase (Thermo Fisher Scientific). Second-strand synthesis was performed with an oligo (dT) anchor primer. The obtained double-stranded DNA was amplified with the anchor-1 primer and nested gene-specific primer SP3_*Bgl*II *nbdA*. The resulting PCR product was purified (MinElute kit, Qiagen) and cloned in pDrive using the Qiagen PCR cloning kit (Qiagen). For the determination of the transcriptional start site 10 individual clones were sequenced.

## RESULTS

### Determination of transcriptional start site of *nbdA* reveals a regulatory region with RpoS and AmrZ binding sites

Automated annotation of the genome sequence of PAO1 predicted the open reading frame (ORF) for *nbdA* to code for 783 amino acid residues. Sequence alignments of the translated sequence with proteins containing N-terminal MHYT-domains revealed a long N-terminal extended region for NbdA. This incited us to analyze the gene region in more detail. In close proximity to the annotated start codon no ribosomal binding site (RBS) could be detected. A 5’-RACE PCR experiment revealed the transcriptional start site of *nbdA* 103 nucleotides downstream of the computationally annotated translation start (Fig. 1). The nearest potential translational start site is 170 nucleotides downstream of the previously annotated translation start and possesses a bona fide ribosome binding site. This results in a shorter ORF coding for a 726 amino acid protein whose N-terminus aligns well with the N-termini of other MHYT-domain proteins. With the new defined transcriptional start, the *nbdA* promoter region was analyzed. Conserved binding sites for the alternative sigma factor RpoS (σ^S^) and the transcription factor AmrZ were identified (34, 35).

**Fig. 1:**
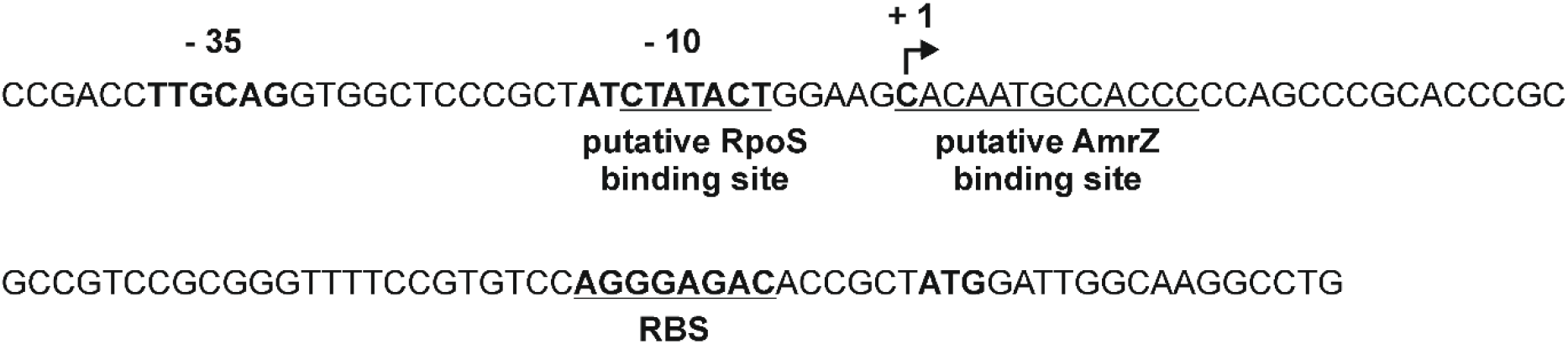
Reannotation of the promoter region of *nbdA* with the experimentally determined transcriptional start site. 5’-RACE PCR experiments revealed transcriptional start site of the *nbdA* gene with a C (+1), 171 nucleotides downstream from the previously annotated ORF start site (Pseudomonas database (36)). The promoter region of *nbdA* was reanalyzed and reveals deduced binding sequences for the alternative σ-factor RpoS and the transcriptional regulator AmrZ.

The binding motif for RpoS lies in the −10 region of the *nbdA* promoter. The predicted AmrZ binding sequence covers the transcriptional start site of *nbdA* indicating a repressor function. In order to investigate the role of the alternative sigma factor RpoS and the transcription factor AmrZ on *nbdA* expression, the promoter region of *nbdA* was transcriptionally fused to the reporter gene *lacZ* and integrated in the φCTX attachment site of PAO1 wt, Δ*rpoS* and Δ*amrZ*. Activity of the β-galactosidase in the respective strains was determined in exponential (4 h) and early stationary (7 h) growth phase. In the wt strain, a 4-fold increase in *nbdA* transcription was observed when cells entered the early stationary phase, which suggests transcriptional activation by RpoS. Deletion of *rpoS* resulted in a loss of *nbdA* promoter activity in both, exponential and stationary growth phase (Fig. 2), confirming the role of RpoS as transcriptional activator of *nbdA*. In the Δ*amrZ* strain a strong increase of *nbdA* promoter activity was observed in both, exponential and early stationary growth phase (Fig. 2). AmrZ is therefore likely acting as a transcriptional repressor for *nbdA*.

**Fig. 2:**
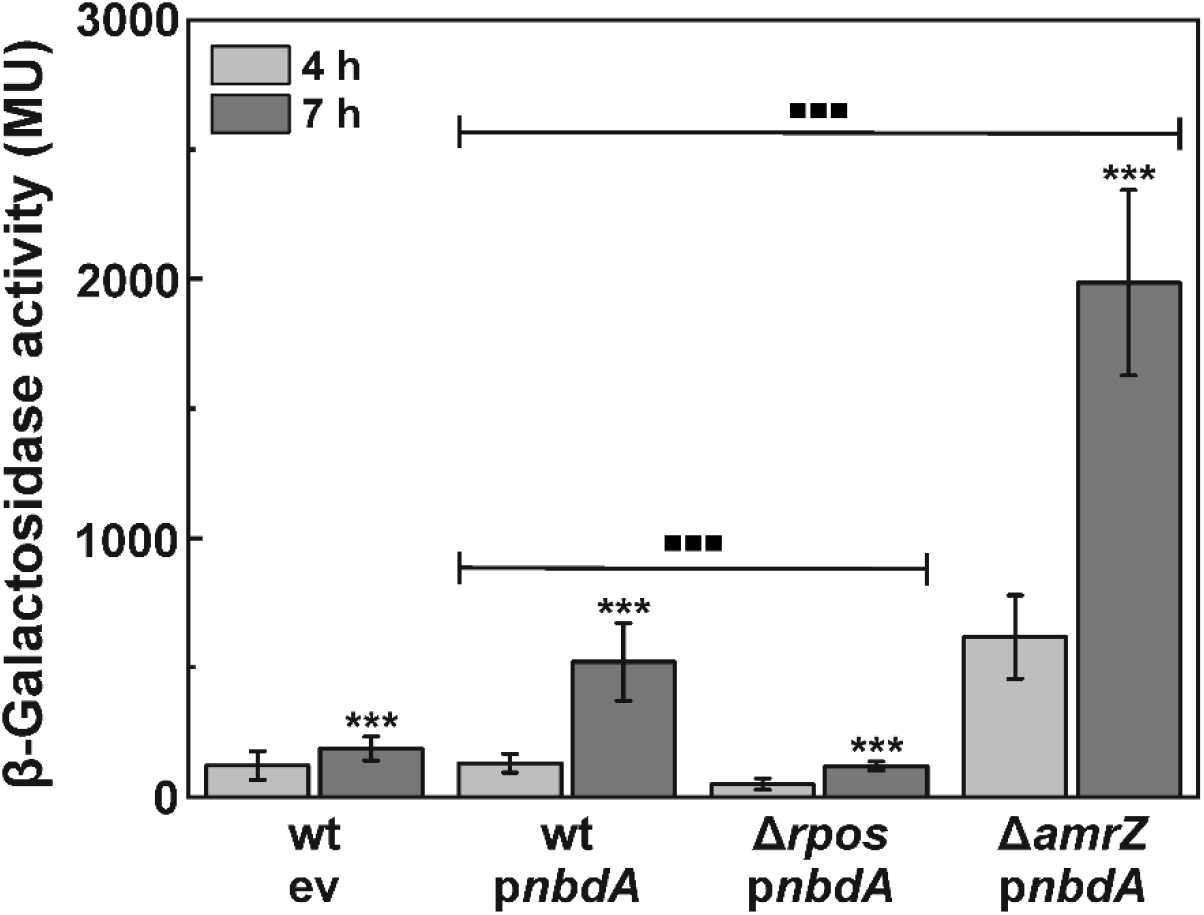
Transcription of *nbdA* is activated by RpoS and repressed by AmrZ. Promoter activity was determined with integrated *pnbdA-lacZ* fusions in late exponential (4 h) and early stationary (7 h) growth phase in wild type (wt) and deletion mutants Δ*rpoS* and Δ*amrZ*. Additionally, the wt with integrated empty vector (ev) was tested for background β-galactosidase activity. The assay was performed in triplicates. Significant changes between 4 and 7 h samples are marked with *. Significant changes in *nbdA* expression levels between deletion mutant strains and the corresponding wt sample are marked with ■ (*** P < 0.001, determined by Student’s T-test).

As there is a sharp oxygen gradient present in biofilm macrocolonies (37), O_2_ might also have an impact on the expression of genes active in biofilms. Although there is no hint for an FNR-like, ANR, or DNR regulator binding site in the promoter region, we tested *nbdA* promoter activity also under anaerobic conditions. Induction of the *nbdA* promoter was observed when cultures reached stationary phase in all tested strains, similarly to the aerobic growth conditions (Fig. 3). Overall, the values for promoter activity under oxygen limitation were significantly lower than in aerobic conditions. The *nbdA* promoter activity in the *amrZ* deletion strain was significantly increased compared to the wild-type background. Therefore, AmrZ seems to repress *nbdA* transcription similarly in aerobic and anaerobic growth conditions.

**Fig. 3:**
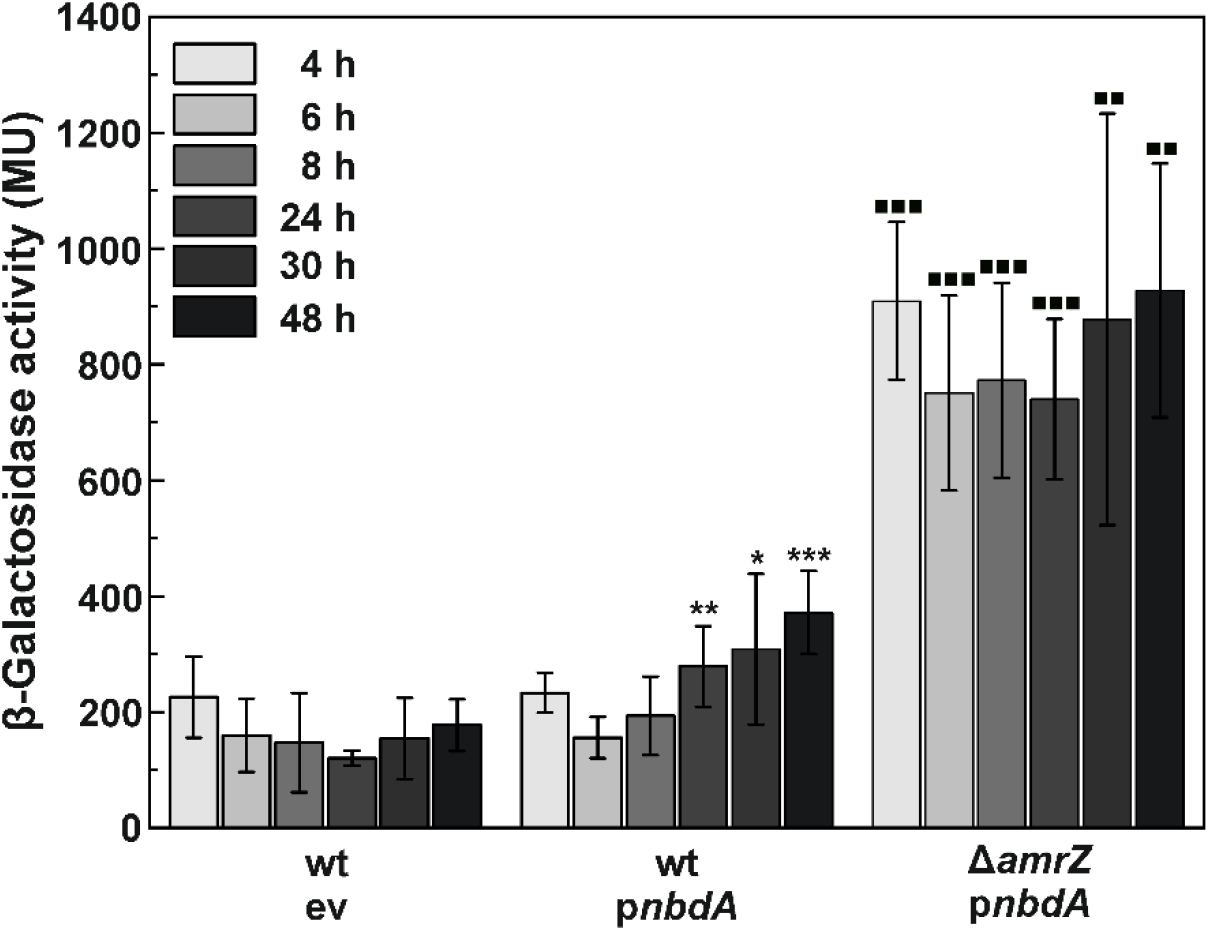
Under oxygen limitation, transcription levels of *nbdA* are induced in stationary growth phase. PAO1 wt and Δ*amrZ* containing an integrated p*nbdA*-*lacZ* fusion and the empty vector control (ev) were grown anaerobically for 48 h in LB medium with 50 mM NaNO3. Inoculation of cultures was performed aerobically therefore the first hours of growth were required to consume remaining oxygen. Expression levels were determined by β-galactosidase assays in triplicates. Significant changes in *nbdA* transcription levels of wt samples to the 6 h sample are marked with **\***. Significant changes in *nbdA* transcription level of Δ*amrZ* samples to corresponding wt levels are marked with ■. (* P < 0.05; ** P < 0.01; *** P < 0.001, determined by Student’s T-test).

### Impact of nitric oxide on the transcription of *nbdA*

We previously reported increased amounts of *nbdA* transcript in dispersed cells after NO-induced biofilm dispersal compared to untreated planktonic cells and suggested a NO-dependent transcriptional regulation of *nbdA* (23). In the light of divergent results on the role of *nbdA* in NO-induced biofilm dispersion (23, 26) we wanted to clarify the regulation of *nbdA* in response to endogenous and exogenous NO. During infections, host macrophages release exogenous NO in order to eradicate bacteria (38). However, under anaerobic conditions, *P. aeruginosa* is able to form endogenous NO during denitrification. Within the denitrification process, nitrite is reduced by the nitrite reductase NirS into NO, which is then further reduced to nitrous oxide by NorCB (39, 40). For its enzymatic activity, NirS requires the incorporation of a heme d_1_ cofactor which is synthesized by NirF (41). Interruption of the denitrification pathway by deletion of the *norCB* gene leads to the accumulation of intrinsic NO under denitrifying conditions (4). A Δ*norCB* strain containing the *nbdA* promoter *lacZ-* fusion was used to analyze the effect of endogenous NO on *nbdA* transcription. Deletion mutants of *nirS* and *nirF*, both unable to form endogenous NO, served as negative controls. The denitrification deficient strains showed normal growth under aerobic conditions in LB medium complemented with KNO_3_ (Fig. 4A). In contrast, under anaerobic denitrifying conditions the growth of PAO1 Δ*nirS* and Δ*nirF* was reduced compared to the wt PAO1 (Fig. 4B). The Δ*norCB* strain was no longer able to grow. For the analysis of *nbdA* transcription, the strains containing the *nbdA* promoter *lacZ*-fusion were grown under aerobic/microaerobic conditions and ß-galactosidase assays were performed with samples of the exponential (4 h) and early stationary growth phase (7 h) (Fig. 4C). Compared to the wt, the *nirS* deletion had a severe activating effect on *nbdA* expression in both, exponential and early stationary growth phase. Surprisingly, *nbdA* transcription in the Δ*nirF* strain, which produces an enzymatically inactive NirS, was not as high as in the Δ*nirS* strain but comparable to the level of transcription in the wt background. The transcription of *nbdA* in the Δ*norCB* strain is slightly decreased compared to the wt background. Due to the impaired growth in anaerobic conditions of Δ*norCB* strain, we could not test for the effect of accumulation of endogenous NO on *nbdA* transcription. In order to confirm the findings for the Δ*nirS* strain, a semi quantitative RT-PCR experiment was performed with cDNA of wt and deletion mutant in both tested growth phases (Fig. 4D). While the control PCR with *recA* primers showed equally strong bands for all samples, *nbdA* expression in the wt in exponential growth phase was weaker than in early stationary growth phase. In the *nirS* deletion strain, there was more transcript of *nbdA* detectable than in wt, which is consistent to the findings of the β-galactosidase assay.

**Fig. 4:**
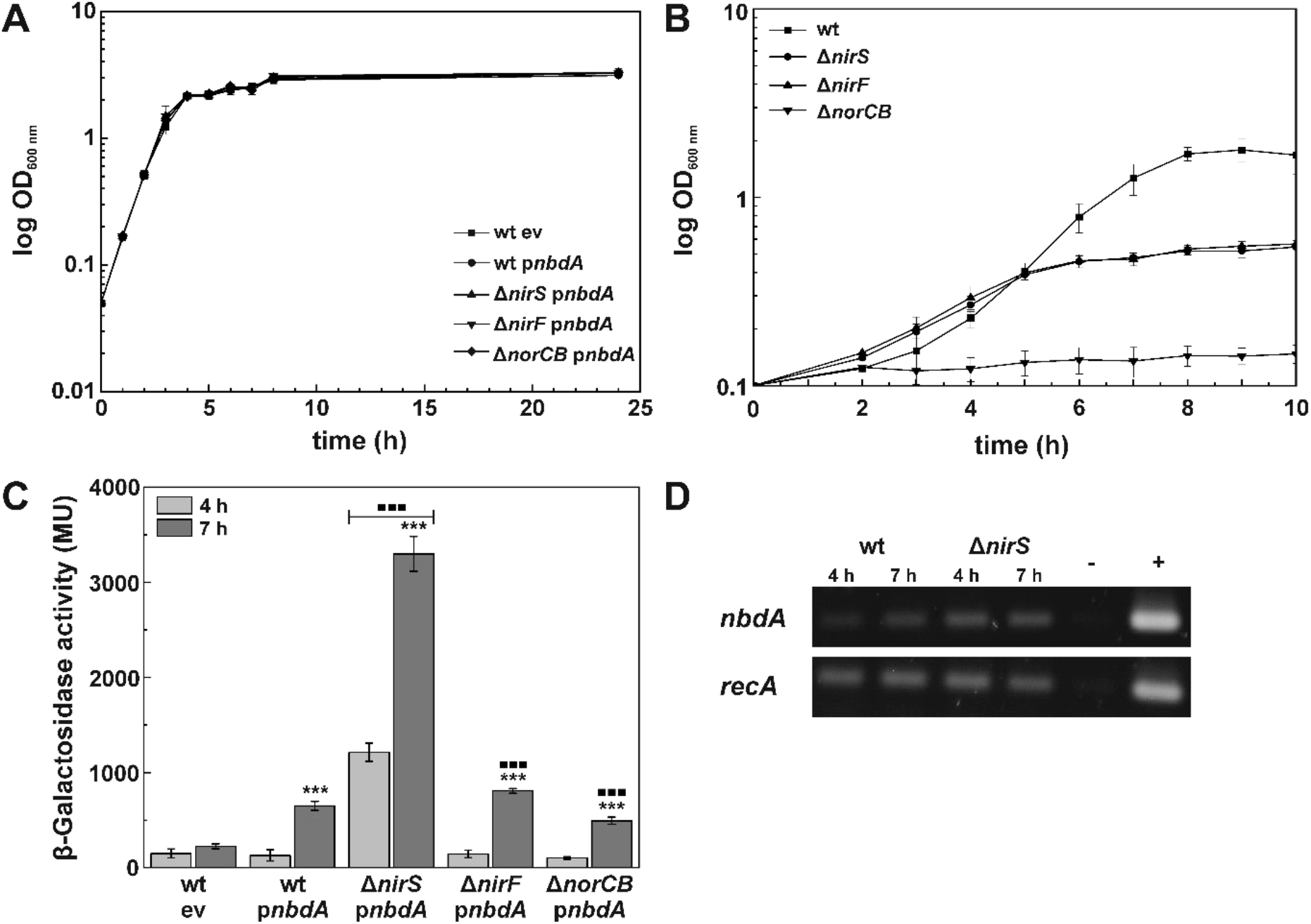
*P. aeruginosa* wild type (wt), Δ*nirS, ΔnirF*, and Δ*norCB* strains harboring a transcriptional *pnbdA lacZ*-fusion and the wt containing the “empty vector” control (ev) were grown in LB supplemented with 50 mM KNO_3_. [**A**] Growth in glass culture flasks was monitored during 24 h in three biological replicates. [**B**] Anaerobic growth in LB medium supplemented with 50 mM KNO_3_ of the wt and deletion mutants was analyzed in sealed bottles for 10 h in biological triplicates. [**C**] β-galactosidase activity was determined from cells aerobically/ microaerobically grown to exponential (4 h) and early stationary phase (7 h). All assays were performed in triplicates. Significant changes between 4 and 7 h samples are marked with *. Significant changes in *nbdA* expression levels between deletion mutant strains and the corresponding wt sample are marked with ■. (*** P < 0.001, determined by Student’s T-test). [**D**] Semi quantitative RT-PCR for *nbdA* transcript and the control *recA* was performed. RNA was extracted from wt and Δ*nirS* strains after 4 and 7 h growth in LB medium.

In addition to the influence of intrinsic nitric oxide on *nbdA* transcription, the effect of exogenous NO was investigated. Therefore, the PAO1 wt harboring the *nbdA-lacZ* fusion was grown with increasing amounts of the NO donor sodium nitroprusside (SNP) and *nbdA* promoter activity was determined by β-galactosidase activity (Fig. 5A). Low concentration of added SNP to the growth medium had no effect on *nbdA* promoter activity, whereas the addition of 500 μM SNP led to a decrease of *nbdA* transcription. This effect is comparable to the observed decrease of *ndbA* transcription in the NO-accumulating strain Δ*norCB*. As *P. aeruginosa* is able to detoxify nitric oxide via flavohemoglobin (42, 43) under aerobic conditions, the influence of short-term NO stress on *nbdA* transcription was analyzed.

**Fig. 5:**
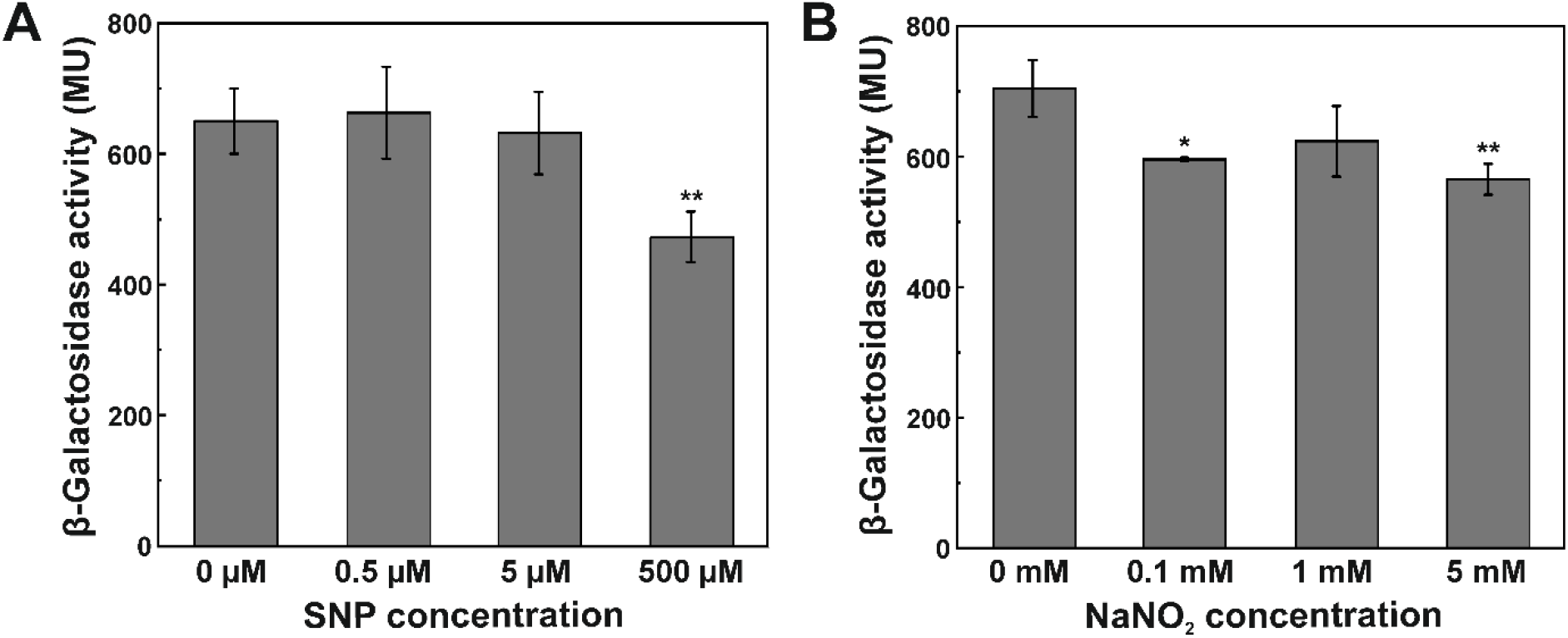
Analysis of transcriptional activity of *nbdA* promoter in response to exogenic NO and NO_2_ sources. Reporter strain PAO1 wt::p*nbdA*-*lacZ* was grown to early stationary phase (7 h) in medium containing increasing amounts of the NO-releasing compound sodium nitroprusside (SNP) [**A**] or NaNO_2_ [**B**]. β-galactosidase activity was measured in triplicates (* P < 0.05; ** P < 0.01, determined by Student’s T-test).

Therefore, PAO1 wt *nbdA-lacZ* was grown to stationary phase in LB and then stressed for 30 min by the addition of 500 μM SNP. Compared to the untreated control, no changes in the *nbdA* promoter activity were observed (data not shown).

In order to figure out whether the strong increase of *nbdA* expression in the Δ*nirS* strain was based on nitrite accumulation due to interrupted denitrification (4), β-galactosidase assays were performed with different amounts of nitrite in the growth medium (Fig. 5B). None of the tested nitrite concentrations had a comparable effect on the *nbdA* promoter as the deletion of *nirS*. The addition of nitrite to the medium rather decreased expression of *nbdA* slightly, probably due to bacteriostatic effect of nitrite.

### DISCUSSION

In this study we analyzed the transcriptional regulation of *nbdA* coding for the phosphodiesterase NbdA, involved in the c-di-GMP modulating network in *P. aeruginosa*. Determination of the transcription initiation site of *nbdA* by 5’-RACE revealed an erroneous annotation of the ORF in the databases. A new promoter region was identified, containing putative binding sites for RpoS and AmrZ. Gene expression of *nbdA* was shown to be activated in stationary growth phase by the alternative sigma factor RpoS (σ^S^). A further level of regulation is introduced through the repression by the ribbon-helix-helix transcription factor AmrZ. Oxygen limitation, supplementation with nitrite, and endogenous or exogenous nitric oxide did not affect the transcription of *nbdA*. Surprisingly, deletion of the nitrite reductase NirS showed a strong activating effect on *nbdA* transcription, while a strain with an enzymatically inactive NirS (Δ*nirF*) showed no transcriptional changes.

The sigma factor RpoS is known as the master regulator of gene expression during stationary growth phase. Furthermore, it is responsible for the activation of genes in response to different stresses, e.g. starvation, heat, oxygen or osmotic stress (44–46). In some Proteobacteria, RpoS is additionally involved in regulation of virulence genes, quorum sensing and motility (47–50). In *E. coli*, RpoS has been shown to play an important role in biofilm maturation, architecture and density (51–53). This link is partly due to the involvement of RpoS in the c-di-GMP regulatory network of *E. coli*. In the *E. coli* K12 strains MC4100 and W3110 a great subset of GGDEF/EAL-domain encoding genes was identified to be under control of RpoS (54, 55). Similarly, in *Pseudomonas* sp. several genes related to biofilm formation, maturation and architecture were shown to be regulated by RpoS (56, 57). A global analysis of *P. aeruginosa* PA14 revealed that 30 out of 40 genes encoding for c-di-GMP modulating enzymes, are either transcriptionally activated or repressed by RpoS (Table 3 and references therein). Thus, similar to *E. coli*, RpoS-dependent regulation significantly affects the c-di-GMP network of *P. aeruginosa*. RpoS regulated genes are often subject to further regulatory mechanisms. Activator or repressor proteins might be involved, as well as post-transcriptional regulation. The *psl* operon coding for matrix polysaccharide biosynthesis genes in *P. aeruginosa* is controlled transcriptionally by RpoS and post-transcriptionally by RsmA (58). In *P. putida* KT2440 the exopolysaccharide cluster *pea* is activated by RpoS and repressed by AmrZ (59). Actually, when we evaluated and compared the data of the PA14 RpoS regulon (35) and the PAO1 AmrZ regulon (34) we found 18 out of 40 genes encoding for c-di-GMP modulating enzymes in *P. aeruginosa* presumably regulated by both proteins, RpoS and AmrZ (Table 3, (34, 35)).

**Table 3:** Genes coding for c-di-GMP modulating proteins in PAO1 and PA14 and their association to RpoS or AmrZ regulon. Data extracted from Schulz 2015, and Jones 2014. RpoS regulon was analyzed in *P. aeruginosa* PA14 via mRNA profiling of Δ*rpoS* vs. wt (35). AmrZ regulon was measured via RNA-Seq experiments in a PAO1 *amrZ* complementation strain vs. Δ*amrZ* (34). GGDEF: diguanylate cyclase domain, EAL: phosphodiesterase domain, GGDEF-EAL: tandem diguanylate cyclase – phosphodiesterase domain, HD-GYP phosphodiesterase domain. + activation; - repression; none: is not part of the indicated regulon.

| PAO1<br>gene locus | PA14<br>gene locus | name | domain(s) | RpoS regulation | AmrZ regulation |
| --- | --- | --- | --- | --- | --- |
| PA0169 | PA14_02110 | *siaD* | GGDEF | - | - |
| PA0285 | PA14_03720 |  | GGDEF-EAL | none | - |
| PA0290 | PA14_03790 |  | GGDEF | none | none |
| PA0338 | PA14_04420 |  | GGDEF | - | - |
| PA0575 | PA14_07500 | *rmcA* | GGDEF-EAL | + | none |
| PA0847 | PA14_53310 |  | GGDEF | + | none |
| PA0861 | PA14_53140 | *rbdA* | GGDEF-EAL | + | none |
| PA1107 | PA14_50060 | *roeA* | GGDEF | + | - |
| PA1120 | PA14_49890 | *tpbB* | GGDEF | + | none |
| PA1181 | PA14_49160 |  | GGDEF-EAL | + | + |
| PA1433 | PA14_45930 |  | GGDEF-EAL | + | none |
| PA1727 | PA14_42220 | *mucR* | GGDEF-EAL | + | - |
| PA1851 | PA14_40570 |  | GGDEF | + | none |
| PA2072 | PA14_37690 |  | GGDEF-EAL | + | + |
| PA2133 | PA14_36990 |  | EAL | + | none |
| PA2200 | PA14_36260 |  | EAL | + | none |
| PA2567 | PA14_31330 |  | GGDEF-EAL | none | - |
| PA2572 | PA14_30830 |  | HD-GYP | + | + |
| PA2870 | PA14_26970 |  | GGDEF | + | + |
| PA3177 | PA14_23130 |  | GGDEF | none | + |
| PA3258 | PA14_21870 |  | GGDEF-EAL | none | none |
| **PA3311** | **PA14_21190** | ***nbdA*** | **GGDEF-EAL** | **+** | **-** |
| PA3343 | PA14_20820 | *hsbD* | GGDEF | + | - |
| PA3702 | PA14_16500 | *wspR* | GGDEF | + | + |
| PA3825 | PA14_14530 |  | EAL | + | + |
| PA3947 | PA14_12810 | *rocR* | EAL | + | none |
| PA4108 | PA14_10820 |  | HD-GYP | + | none |
| PA4332 | PA14_56280 | *sadC* | GGDEF | none | - |
| PA4367 | PA14_56790 | *bifA* | GGDEF-EAL | + | - |
| PA4396 | PA14_57140 |  | GGDEF | none | none |
| PA4601 | PA14_60870 | *morA* | GGDEF-EAL | + | - |
| PA4781 | PA14_63210 |  | HD-GYP | + | + |
| PA4843 | PA14_64050 | *gcbA* | GGDEF | none | + |
| PA4929 | PA14_65090 |  | GGDEF | + | - |
| PA4959 | PA14_65540 | *fimX* | GGDEF-EAL | + | + |
| PA5017 | PA14_66320 | *dipA* | GGDEF-EAL | none | - |
| PA5295 | PA14_69900 | *proE* | GGDEF-EAL | + | none |
| PA5442 | PA14_71850 |  | GGDEF-EAL | - | - |
| PA5487 | PA14_72420 | *dgcH* | GGDEF | + | none |
| * | PA14_59790 | *pvrR* | EAL | none | none |

\* gene not present in PAO1

The transcriptional regulator AmrZ controls a large regulon containing 398 gene regions in PAO1. Transcription of *amrZ* itself is in a great extend dependent on the alternative sigma factor AlgT (σ^22^) which is known to regulate coversion to mucoidity and stress responses in *P. aeruginosa* (60, 61). AmrZ was shown to regulate genes important for *P. aeruginosa* virulence, including type IV pili, extracellular polysaccharides, and the flagellum (34). It particularly influences genes required for alginate production and twitching-motility (34, 62–64). Within the c-di-GMP network of *P. aeruginosa*, AmrZ activates transcription of 14 genes and represses 10 genes encoding GGDEF/EAL-domain proteins ((34), Table 3). With these numbers, AmrZ appears to be one of the major regulators for genes coding for c-di-GMP modulating enzymes in *P. aeruginosa*, possibly affecting the cellular c-di-GMP level. This role for AmrZ was previously also observed in *P. fluorescens* F113, where the cellular c-di-GMP level was affected by AmrZ through the regulation of a complex network of genes encoding DGCs and PDEs (65). From our work we conclude that *nbdA* transcription is repressed by AmrZ during aerobic as well as anaerobic planktonic growth while a condition in which the *nbdA* promoter is de-repressed remains uncertain. Repression through AmrZ is described to be dependent on the C-terminus mediated tetramerization of the protein (66). In some cases, e.g. *pilA* repression, the expression level of AmrZ plays an important role for its function, as binding efficiency of AmrZ to different promoter regions differs (64). Additionally, a competition of the activator RpoS and the repressor AmrZ upon binding to the *nbdA* promoter might be possible.

### Effects of endogenous or exogenous nitric oxide on *nbdA* expression

In our previous study we observed elevated transcription levels of *nbdA* in cells dispersed from biofilms after NO-treatment when compared to planktonically grown cells in RT-qPCR experiments (23). Therefore, we suggested transcriptional regulation of *nbdA* by NO in this biofilm model. For *P. aeruginosa* two NO-responsive transcriptional regulators, FhpR and DNR are described. DNR is a heme-containing CRP/FNR type regulator that specifically activates denitrification genes under anaerobic conditions (67, 68). Transcription of the second NO responsive regulator in *P. aeruginosa*, FhpR, is σ^54^-dependent and activates flavohemoglobin expression under aerobic conditions for the detoxification of NO in the cell (42, 43). When analyzing the reannotated *nbdA* gene and promoter region, no similarity with either the FhpR or DNR consensus binding site was detected (69). Therefore, a direct influence of NO on *nbdA* transcription was unlikely. These findings in addition to the contradictory results in the literature concerning the involvement of NbdA in NO-induced biofilm dispersal of *P. aeruginosa* (23, 26) led to the reevaluation of the transcriptional regulation of *nbdA* by NO. In this study, no direct stimulation of *nbdA* promoter activity by NO, neither by addition of exogenous NO nor by accumulation of intrinsic NO in planktonically grown cells was observed. The previously observed induction of *nbdA* expression in our qRT-PCR experiments (23) might be due to more complex regulatory processes during biofilm formation and dispersal. From the present data, we conclude that *nbdA* expression in planktonic cells is not directly induced by NO at the transcriptional level.

### Effect of the nitrite reductase NirS on *nbdA* promoter activity

In this study, we observed a strong increase in the *nbdA* transcription level when the nitrite reductase NirS was deleted. At first, we assumed that the upregulation of *nbdA* expression might be due to accumulation of intrinsic nitrite from interrupted denitrification. However, addition of nitrite to the growth medium did not change *nbdA* promoter activity. Further, the Δ*nirF* strain producing an enzymatically inactive NirS protein (41) did not enhance *nbdA* transcription. Therefore, we suggest that the presence of the periplasmic protein NirS affects *nbdA* transcription independently of its enzymatic activity. The moonlighting role of NirS was previously described for the type III secretion system in *P. aeruginosa* (70). Additionally, NirS was shown to affect flagellum biogenesis by the formation of a complex with the flagellar subunit FliC and the chaperone DnaK (4, 60, 71). Suggesting this complex role for NirS besides denitrification in *P. aeruginosa*, the increase of *nbdA* promoter activity in the Δ*nirS* strain is probably derived from a global regulatory change in the cell.

All in all, we were able to reannotate the *nbdA* gene and revealed consensus sequences for the alternative sigma factor RpoS and the transcription factor AmrZ within the *nbdA* promoter region. Our data confirmed RpoS as activator and AmrZ as repressor for *nbdA* transcription, however, no transcriptional regulation by endogenous or exogenous NO or nitrite was observed in planktonically grown cells.

## Abbreviations

c-di-GMP: bis-3,5-cyclic di-guanosine monophosphate
DGC: diguanylate cyclase
NO: nitric oxide
PDE: phosphodiesterase
RACE: rapid amplification of cDNA ends

## AUTHORS STATEMENTS

### Authors and contributors

KG, SZ, NFD conceived the study, KG and SZ performed experiments, KG and SZ analyzed the data, KG wrote first draft of manuscript, all authors revised and approved the final version of the manuscript.

## Acknowledgements

We thank Sandra Schwarz (Tübingen, Germany) for the generous gift of the plasmid pEXG2.

